# FRAPPe - a novel FRET-based biosensor to detect inorganic polyphosphates

**DOI:** 10.64898/2026.01.30.702422

**Authors:** Sunayana Sarkar, Payal Agrawal, Harsha Sharma, Azmi Khan, Manisha Mallick, Deepti Ranjan Pradhan, Sandra Moser, Sreejith Raran Kurussi, Henning Jessen, Kaustubh R Mote, Vinay Bulusu, Rashna Bhandari, Manish Jaiswal

## Abstract

Inorganic polyphosphate (polyP), a linear polymer of orthophosphate residues, is involved in a range of cellular processes, including phosphate storage, blood clotting, chromatin remodeling, RNA processing, and mitochondrial function. Despite its significance, tools for monitoring polyP with specificity and in real time are lacking, limiting insights into its dynamics. Here, we present FRAPPe (FRET-based Ratiometric Analysis of Polyphosphate), a FRET-based sensor for polyP. FRAPPe consists of an eCFP–eYFP FRET pair flanking CHAD, a polyP-binding domain. The sensor remains unresponsive to nucleic acids and other polyanions, while polyP binding induces a significant decrease in FRET, allowing quantitative estimation of polyP. We further show that FRAPPe can qualitatively estimate polyP levels from crude cell extracts, supporting its utility in high-throughput genetic and pharmacological screens to identify regulators of polyP metabolism and signaling.

## Introduction

Inorganic polyphosphate (polyP), a linear polymer of orthophosphate residues linked by phosphoanhydride bonds, is conserved from bacteria to humans (1–5). Recently, PolyP has been implicated in stress responses (e.g., starvation and oxidative stress adaptation), ionic buffering, blood clotting, wound healing, mitochondrial function, development, and neural function (2,5–19). One of the well-documented functions of polyP in higher eukaryotes is its role in blood clotting, a function that is conserved between flies and humans (5,8,15,16,20,21). Ongoing research continues to discover new biological and molecular functions of polyP in cell biology and pathogenesis, establishing its significance in health and disease.

Over many years, several biochemical and cellular techniques have been developed to detect polyP in various organisms. PolyP levels have also been qualitatively detected using gel electrophoresis followed by Toluidine Blue O (TBO) staining, which gives a purple color upon binding to polyP (22,23). While this method is used in organisms with high polyP content, such as yeast and bacteria, it is not particularly sensitive in higher eukaryotes due to their lower polyP content (23). DAPI-based detection is one of the most widely used methods to quantify polyP, as when DAPI binds to polyP, its emission spectra show a characteristic red shift with a maxima at 550 nm on excitation at 415 nm DAPI-based detection, however, can be severely misinterpreted in the presence of RNA in the sample due to the overlapping emission spectral characteristics of DAPI-PolyP and DAPI-RNA, particularly when polyP levels are very low (24,25). The widely used assay for *in vitro* detection of polyP is based on its enzymatic digestion with polyphosphatases, followed by measurement of liberated monophosphates (Pi) using malachite green or ascorbic acid (26,27). To detect polyP *in situ,* DAPI and JCD8 dyes have been used. In addition, an epitope-tagged polyP-binding domain (PPBD) is used for in situ polyP detection (28). Limitations of existing methods include the inability to study polyP dynamics in live cells, the need for tedious procedures, and their inability to support high-throughput screening to identify polyP regulators.

In this work, we developed FRAPPe (FRET-based Ratiometric Analysis of Polyphosphate), a Förster Resonance Energy Transfer (FRET)- based biosensor that enables ratiometric detection of polyP binding via conformational changes.

## Results

### Design of the polyP sensor- FRAPPe

We designed the FRAPPe construct by assembling, in sequence, an N-terminal polyhistidine tag, a FLAG tag, the donor fluorophore eCFP (with nine C-terminal residues truncated), a three-amino acid linker, the polyP-binding domain ssCHAD, a four-amino acid linker, and the acceptor fluorophore eYFP (lacking its first two N-terminal residues). In **Fig. 1A**, we show the schematic of this construct along with its AlphaFold 3.0-predicted structure. The predicted model indicates that in the absence of polyP, the eCFP and eYFP chromophores are separated by 40.3 Å, a distance that supports efficient FRET. When we docked polyP-6 into the CHAD binding pocket, AlphaFold predicted a 1.3 Å increase in the interchromophore distance, suggesting a potential reduction in FRET efficiency upon polyP binding.

**Fig. 1:**
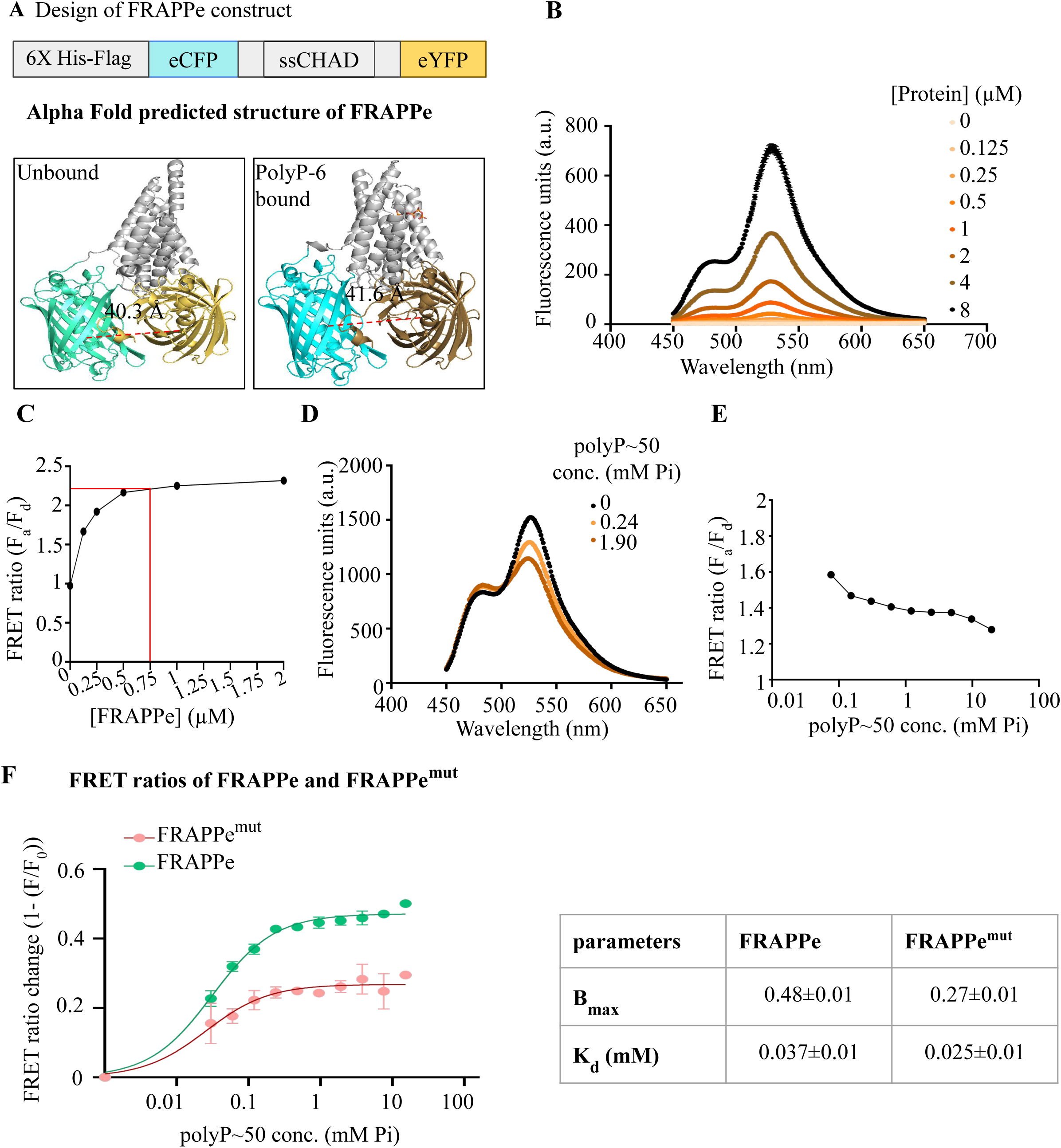
FRAPPe, a FRET based sensor for polyP. **A)** Schematic of the FRAPPe construct, and Alpha Fold 3.0 predicted model of FRAPPe (unbound) showing distance between two chromocentres at 40.3 Å, and FRAPPe bound to polyP-6 (docked) showing distance between two chromocentres at 41.6 Å. **B)** Fluorescence spectral scan (450nm-650nm) of different concentrations of FRAPPe on excitation at 415 nm. **C)** FRET ratio calculated from *1B* showing 0.75 µM in red lines, after which FRET ratio does not vary with changing FRAPPe concentration. FRAPPe concentration chosen for further experiments is 0.75 µM. **D)** Fluorescence spectral scan (450nm-650nm) of 0.75 µM FRAPPe on excitation at 415 nm with varying concentrations of commercially available polyP∼50. **E)** FRET ratio calculated from *1D*. **F)** FRET ratio change of FRAPPe and FRAPPe^MUT^ across polyP∼50 concentration range 0-30.6mM (Pi terms). The chart contains the binding affinity (K_d_) and the binding capacity (B_max_) of the wild-type and mutant sensors. Data plotted with s.e.m. as an error bar.

We cloned the FRAPPe construct into the pRSET vector backbone and expressed it in *E. coli*. We purified three independent batches of FRAPPe protein using affinity chromatography followed by size-exclusion chromatography **(see Materials and Methods)**. To determine the optimal sensor concentration for FRET measurements, we recorded emission spectra (excitation: 415 nm; emission: 450–550 nm) across a range of protein dilutions. We then calculated FRET ratios (Fa/Fd) by dividing the average eYFP emission (515–525 nm) by the average eCFP emission (480–490 nm). The FRET ratio plateaued above 1 μM protein concentration **(Fig. 1B–C)**, indicating saturation. Based on this, we selected 0.75 μM FRAPPe for all subsequent assays.

To test whether polyP binding alters FRET in FRAPPe, we recorded fluorescence emission spectra (excitation: 415 nm; emission: 450–550 nm) in the presence of increasing concentrations of polyP (∼50-mers). As polyP levels increased, we observed a progressive rise in donor (eCFP) emission and a corresponding drop in acceptor (eYFP) emission **(Fig. 1D, Supplementary Fig. 1A)**. This shift indicates a reduction in FRET upon polyP binding. These results align with our structural predictions, which suggest that polyP binding increases the distance between the fluorophores in the FRAPPe sensor.

We analyzed the FRET ratio (Fa/Fd) across increasing concentrations of polyP using data from the spectral scans **(see Materials and Methods)**. The FRET ratio decreased sharply and reached an initial plateau within the 0–15 mM polyP range (expressed in Pi units), followed by a second phase of gradual decline that plateaued at concentrations as high as 977.5 mM **(Supplementary Fig. 1B)**. This biphasic response suggests potential two-site binding kinetics. However, since the second phase occurs at polyP concentrations far exceeding physiological levels reported in any organism to date, we focused our analysis on the first phase using a one-site binding model. **Fig. 1E** shows the FRET response curve for polyP concentrations up to the first plateau.

To investigate the basis of the two-site binding behavior observed in the FRET measurements, we performed isothermal titration calorimetry (ITC) to assess the binding kinetics between FRAPPe and polyP (PolyP-100) (see Materials and Methods). The ITC results confirmed that FRAPPe binds polyP with an affinity consistent with that inferred from the FRET data. However, the binding affinity of FRAPPe (∼5 μM) was lower than that of the standalone CHAD domain, which has been reported to bind polyP with ∼30 nM affinity based on Grating Coupled Interferometry (GCI) measurements by Lorenzo-Ortiz and colleagues (29). The representative ITC curve further indicated that FRAPPe possesses more than one polyP-binding site, with a stoichiometry of N = 1.42 ± 0.02 **(Supplementary Fig. 1C)**.

To validate the polyP-binding specificity of FRAPPe, we generated a mutant version (FRAPPe^Mut^) by incorporating a CHAD mutant (CHAD^Mut^) in which all residues previously reported to mediate polyP binding were substituted with alanine (29) **(Supplementary Fig. 2A)**. AlphaFold 3.0 structural predictions revealed a root-mean-square deviation (r.m.s.d.) of 0.381 between CHAD and CHAD^Mut^, indicating no significant structural perturbation. Docking simulations with polyP-6 predicted a complete loss of polyP binding in CHAD^Mut^ **(Supplementary Fig. 2B)**.

To experimentally validate this, we compared polyP binding between GST-CHAD and GST-CHAD^Mut^ using a Bis-biotin-polyP8 pull-down assay **(Supplementary Fig. 2C–D)**. We observed a marked reduction in polyP binding to GST-CHAD^Mut^, with its binding signal indistinguishable from GST alone **(Supplementary Fig. 2E)**. These results confirm that CHAD^Mut^ is unable to bind polyP and validate the specificity of the wild-type CHAD domain in FRAPPe.

We performed AlphaFold 3.0-based structural predictions for both FRAPPe and FRAPPe^Mut^ and observed an r.m.s.d. of 2.438, primarily reflecting minor positional shifts in the C-terminal His and FLAG tags, with no significant alterations in the core structure **(Supplementary Fig. 2G–H)**. To compare their functional responses, we analyzed FRET changes in FRAPPe and FRAPPe^Mut^ as polyP concentration increased. FRAPPe^Mut^ exhibited a markedly reduced FRET response relative to the wild-type sensor **(Fig. 1F)**, consistent with impaired polyP binding. Together, these results confirm that FRAPPe responds specifically to polyP, and that this interaction is responsible for the observed FRET reduction.

### FRAPPe specifically detects polyP over other negatively charged polymers

To evaluate the specificity of FRAPPe for polyP, we tested its response to physiologically relevant concentrations of negatively charged polymers, namely, DNA, RNA, and heparin (30,31). These polymers did not alter the FRET signal of polyP-unbound FRAPPe **(Fig. 2A)**, indicating that the sensor does not bind nonspecifically to other polyanions. Moreover, upon titrating polyP∼50 in the presence of these polymers, we observed no significant change in the sensor’s binding kinetics, including binding affinity (Kd) or maximal binding capacity (Bmax) **(Fig. 2B, 2E)**.

**Fig. 2:**
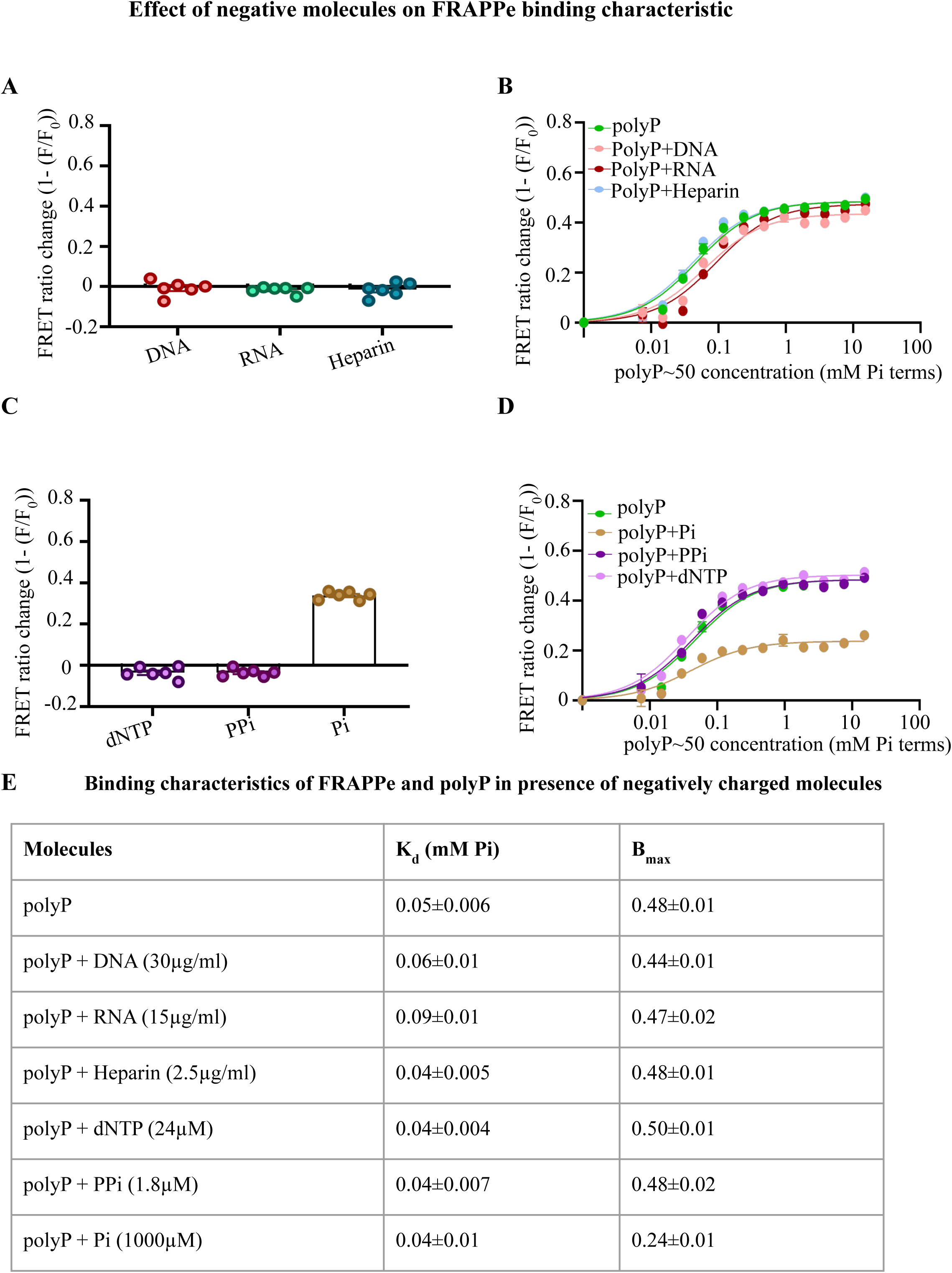
FRAPPe specifically binds to polyP. **A)** FRET ratio change of FRAPPe protein on exposure to negatively charged polymers- DNA (30µg/ml), RNA(15µg/ml), Heparin (2.5µg/ml) in 20 mM Tris buffer. **B)** FRET ratio change of FRAPPe protein across polyP∼50 concentration range 0-30.6mM (Pi terms) in presence of negatively charged polymers- DNA (30µg/ml), RNA(15µg/ml), Heparin (2.5µg/ml) in 20 mM Tris buffer. **C)** FRET ratio change of FRAPPe protein on exposure to negatively charged moieties- dNTP (24µM), PPi(2.18µM), Pi (1mM) in 20 mM Tris buffer. **D)** FRET ratio change of FRAPPe protein across polyP∼50 concentration range 0-30.6mM (Pi terms) in presence of negatively charged moieties- dNTP (24µM), PPi (2.18µM), Pi (1mM) in 20 mM Tris buffer. The chart highlights the binding affinity (K_d_)and the binding capacity (B_max_) of FRAPPe towards polyP in the presence of negatively charged moieties. Data plotted with s.e.m. as an error bar. **E)** Table showing the K_d_ and B_max_ of each of the ligands used in Fig. 2.

Next, we assessed the influence of phosphate-containing small molecules, namely, dNTPs, pyrophosphates, and monophosphates, on FRAPPe’s FRET response. The FRET ratio remained unchanged in the presence of dNTPs and pyrophosphates. However, monophosphates caused a notable decrease in the FRET ratio **(Fig. 2C)**. When polyP∼50 was added to these solutions, the sensor’s affinity for polyP remained unchanged, but the B_max_ was reduced in the presence of monophosphates **(Fig. 2D, 2E)**.

To further investigate this, we tested direct binding of FRAPPe to monophosphates and confirmed that it binds them with a significantly lower affinity than polyP **(Supplementary Fig. 3A)**. These results suggest that while FRAPPe maintains high specificity for polyP, its use may be limited in environments with high monophosphate concentrations. We recommend using phosphate-deficient buffers for optimal performance. This limitation does not apply to experiments using extracted polyP, as the extraction procedure removes monophosphates.

### Validation of FRAPPe using polyP from yeast and mammalian tissue

To assess the utility of FRAPPe in detecting organismal polyP, we first tested its ability to measure polyP extracted from *Saccharomyces cerevisiae*. We compared wild-type (BY4741) cells with a *vtc4Δ* mutant strain known to lack polyP due to defective polyphosphate kinase activity (32–34). A 40 mL culture (OD₆₀₀ ≈ 1.5) was split equally. One-half was used for polyP extraction via the phenol-chloroform method, followed by quantification using both the FRAPPe sensor and the Malachite Green assay. The latter detects monophosphates (Pi) released by enzymatic digestion of purified polyP (5,26). In wild-type, we estimated ∼32 nmol of polyP (in Pi equivalents per OD₆₀₀), whereas the *vtc4Δ* mutant showed negligible levels **(Fig. 3A)**. Correspondingly, FRAPPe recorded a greater FRET ratio shift in wild-type samples and a minimal response in *vtc4Δ*, indicating its ability to detect relative changes in extracted polyP **(Fig. 3B)**.

**Fig. 3:**
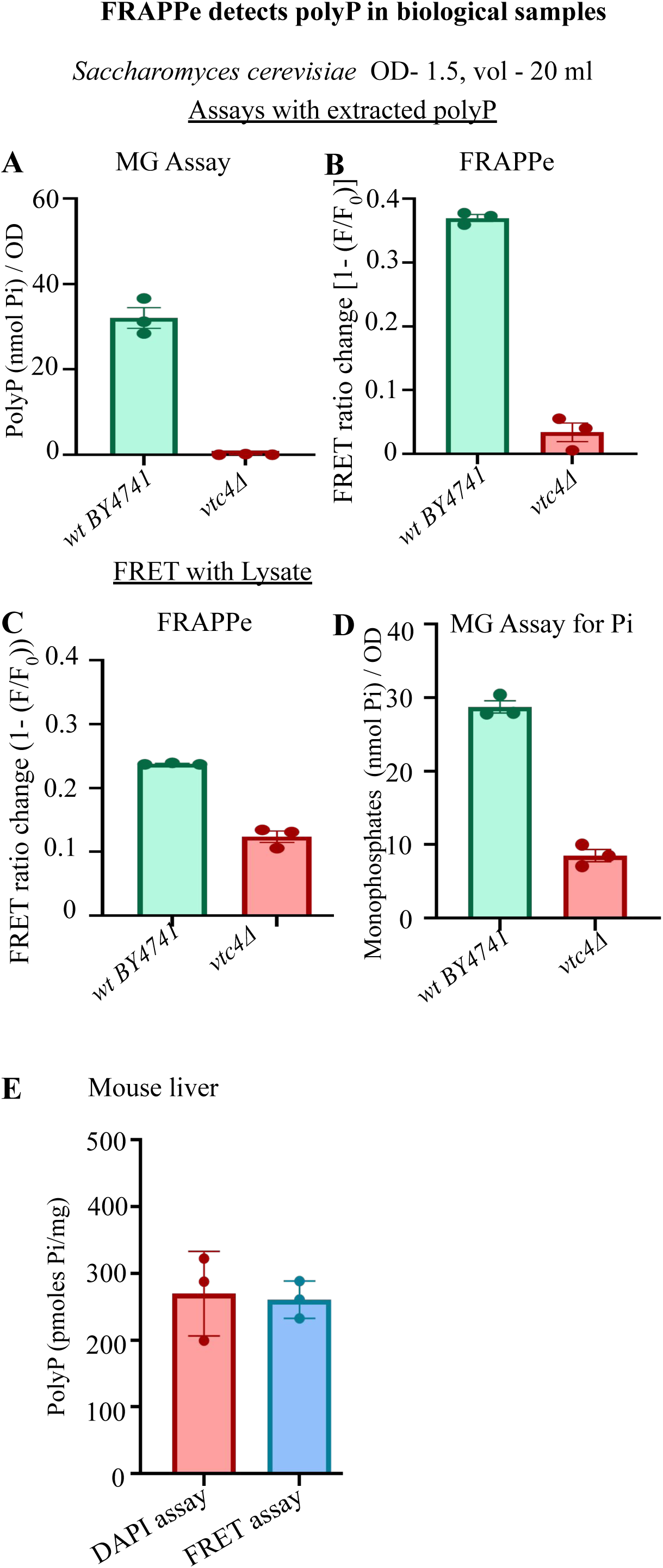
FRAPPe can detect polyP in biological samples. **A-C)** Characterisation of polyP content in *BY4741* (wild-type), and *vtc1Δ* mutants yeast (OD_600_-1.5, vol-10ml) after extraction **(A,B)** and directly from lysate **(C)**, using Malachite Green Assay **(A)** and FRAPPe (**B,C)**. **D)** Characterisation of monophosphate content in *BY4741* wild-type, and *vtc1Δ* mutants (OD_600_-1.5, vol-20ml) during polyP extraction, using Malachite Green assay. **E)** PolyP estimation from mouse liver using DAPI assay and FRAPPe (FRET assay).

Next, we tested FRAPPe on crude lysates from the remaining half of the yeast cultures. Here too, wild-type lysates showed a significantly greater FRET ratio shift than *vtc4Δ* **(Fig. 3C)**. However, unlike extracted polyP samples, *vtc4Δ* lysates still induced a measurable FRET change, which we hypothesized was due to background monophosphates. A comparative Malachite Green assay revealed lower monophosphate levels in *vtc4Δ* than in wild-type **(Fig. 3D)**, supporting the idea that both polyP and monophosphate concentrations influence the FRET signal in crude lysates. Despite this, since the FRET response in *vtc4Δ* serves as a baseline, FRAPPe can still be used for relative polyP quantification in high-throughput genetic or pharmacological screens. These screens should ideally be followed by confirmatory quantification of polyP and Pi from purified extracts.

To extend this utility, we tested FRAPPe on polyP extracted from mouse liver. Absolute quantification of polyP using FRAPPe was comparable to measurements obtained using a DAPI-based biochemical assay **(Fig. 3E)** (36). Together, these results demonstrate that FRAPPe can detect polyP extracted from both yeast and mammalian tissue, supporting its applicability across organisms.

### FRAPPe binding characteristics with variable polyP chain lengths

Given that polyP chain lengths vary across organisms and physiological contexts, we tested whether FRET response is influenced by polymer length. We analyzed sensor performance using polyP-14 (P14), polyP∼50, polyP-65 (P65), and polyP-130 (P130), all at equimolar concentrations (in Pi terms). Two key observations emerged: (1) The apparent binding affinity of FRAPPe decreased with increasing chain length. Specifically, the dissociation constants were: Kd(P14) = 0.048 ± 0.01 mM, Kd(P65) = 0.057 ± 0.01 mM, and Kd(P130) = 0.176 ± 0.1 mM. (2) For longer polymers (P65 and P130), we observed a characteristic bell-shaped response in FRET ratio beyond the first maxima, suggesting a concentration-dependent inhibition-like behavior at higher polymer doses **(Fig. 4A–B)**. Since equimolar phosphate concentrations correspond to different numbers of polymer molecules (more for P14, fewer for P130), we repeated the assay using equimolar polymer concentrations to isolate the effect of polymer copy number. In this setup, we found that FRAPPe binds P65 and P130 with similar high affinity (Kd ≈ 0.0007–0.0008 mM), whereas binding to P14 was weaker (Kd = 0.003 ± 0.0009 mM) **(Fig. 4C)**.

**Fig. 4:**
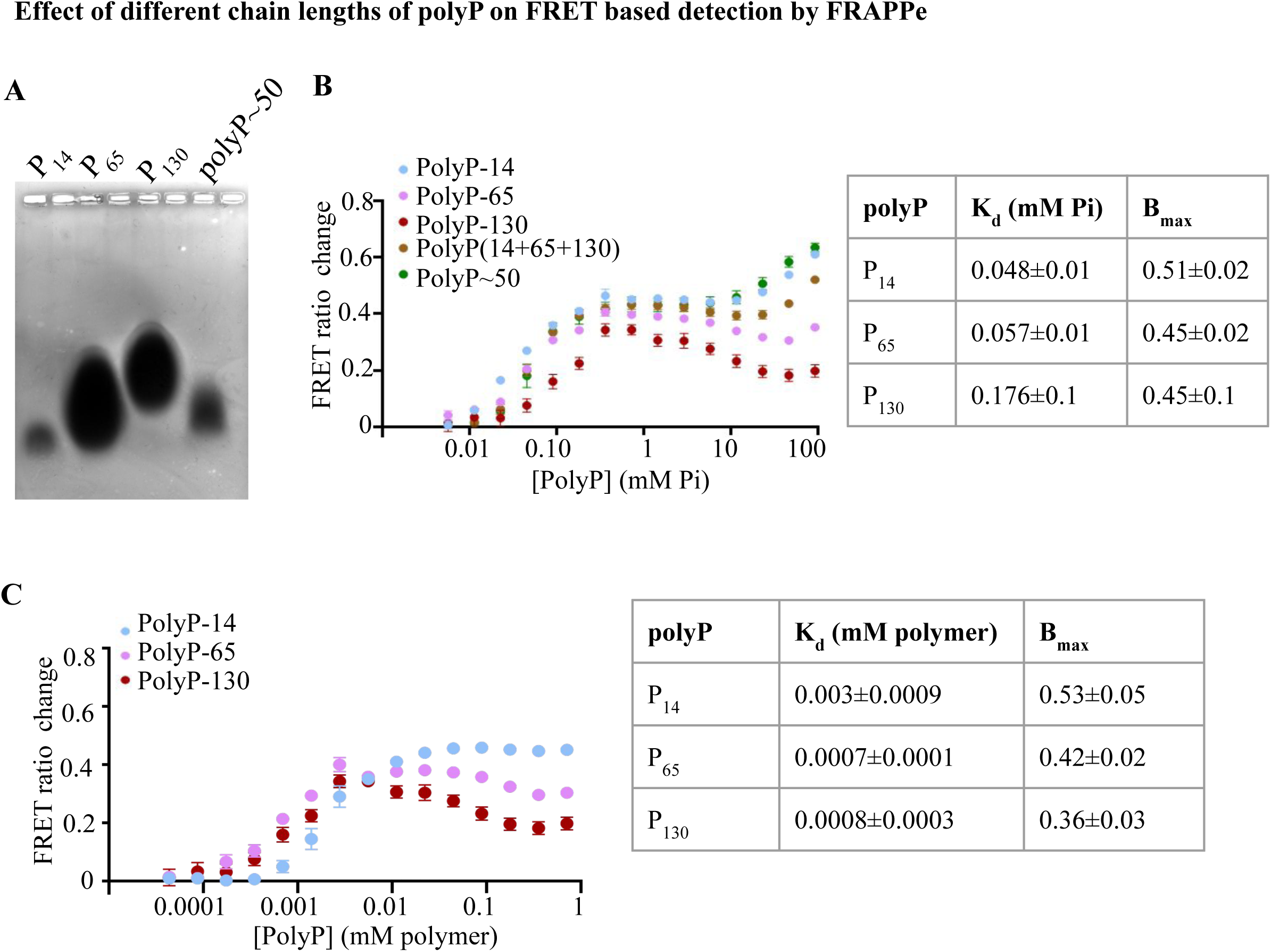
Effect of different chain lengths of polyP on FRET-based detection by FRAPPe. **A)** Agarose gel electrophoresis to show the different chain lengths of polyP used in the experiments - P_14_, P_65_, P_130_, and polyP∼50. **B)** The binding characteristic of FRAPPe to varying chain lengths of polyP, alone and in an equimolar phosphate mixture. The chart represents the binding affinity (K_d_) and the maximum binding capacity (B_max_). **C)** The binding characteristic of FRAPPe to varying chain lengths of polyP with equimolar polymer concentrations. The chart represents the binding affinity (K_d_) and the maximum binding capacity (B_max_).

These results emphasize the need for careful titration when analyzing unknown samples, as both chain length and concentration influence the FRET response. With appropriate standards and interpolations, FRAPPe may also enable estimation of median polyP chain lengths in complex biological samples.

## Discussion

The development of FRAPPe, a sensor for FRET-based Ratiometric Analysis of Polyphosphate, represents a significant advance in the toolkit for studying polyP biology. FRAPPe reliably detects polyP across a wide range of chain lengths and experimental conditions, and exhibits a robust FRET response with high specificity. Notably, the sensor does not respond to other polyanions such as DNA, RNA, or heparin, which have historically confounded biochemical detection of polyP. This selectivity addresses a major limitation of DAPI-based methods, which require stringent controls to eliminate nucleic acid contamination (24,25). By directly binding polyP via the CHAD domain, FRAPPe bypasses enzymatic digestion, eliminating incomplete cleavage or enzyme inhibition and enabling a one-step readout (5,35).

Importantly, FRAPPe demonstrates the capacity to quantify polyP extracted from *Saccharomyces cerevisiae* and mammalian tissue lysates **(Fig. 3)** (36). Its ratiometric readout enables robust quantitative comparisons without the need for RNA digestion or polyP purification (35). This enables its adaptation to plate-based high-throughput screening platforms for the identification of genetic or pharmacological modulators of polyP metabolism and signaling.

We also examined the influence of phosphate-containing small molecules on sensor performance. While FRAPPe remains insensitive to dNTPs and pyrophosphates, it shows reduced efficacy in the presence of monophosphates **(Fig. 2, Supplementary Fig. 3)**. Although the binding affinity (K_d_) for polyP is unaffected by monophosphates, the maximum binding capacity (B_max_) is significantly lowered, suggesting a distinct, non-competitive binding site for Pi on the sensor. This observation warrants further investigation but is currently limited by the absence of a solved structure for the ssCHAD domain and our inability to crystallize the full-length FRAPPe (29). Nevertheless, since polyP extraction protocols eliminate free monophosphates, FRAPPe performs optimally with extracted samples. We recommend using phosphate-deficient buffers (e.g., Tris or HEPES) for lysate-based applications to minimize interference.

We observed that FRAPPe’s response is also influenced by the chain length of polyP, with longer chains producing complex dose-response curves, including apparent inhibition at higher concentrations **(Fig. 4)**. This likely reflects multivalent interactions where multiple sensors engage the same polymer, thereby limiting the extent of FRET reduction. Nonetheless, this behavior provides an opportunity to tune the sensor for different biological contexts or to engineer chain-length preferences in future versions.

In addition to validating FRAPPe against extracted polyP from yeast and liver tissue, we demonstrate its ability to detect polyP directly in crude yeast lysates **(Fig. 3C)**. This allows for rapid, extraction-free screening of polyP phenotypes across genetic backgrounds. Although monophosphate interference is a caveat, its influence can be empirically accounted for by using appropriate controls (34). The insensitivity of FRAPPe to heparin further expands its potential for diagnostic use, where clinical samples may contain diverse polyanions (22).

To our knowledge, FRAPPe is the first genetically encoded FRET sensor for polyP. While this study focuses on *in vitro* validation, its modular architecture enables future *in vivo* adaptation, enabling spatiotemporal monitoring of polyP dynamics in live cells and tissues. Such a capability could catalyze discoveries into the roles of polyP in development, stress signaling, neurobiology, and diseases, areas that remain poorly understood due to the lack of suitable real-time detection tools.

## Material and methods

### Plasmids

cDNA encoding *Ss*CHAD^WT^ procured from Addgene (#127783). cDNA encoding *Ss*CHAD wild-type (CHAD^WT^) and *Ss*CHAD^Mut^ (CHAD^Mut^) were synthesized from Genewiz (pGW-CHAD) and cloned into the pGex-6P-2 vector at EcoRI and XhoI sites using conventional cloning protocols. The constructs have N-terminal glutathione S-transferase (GST) and Xpress tags.

### Expression and purification of GST, GST-*Ss*CHAD^WT^ and GST-*Ss*CHAD^Mut^

BL21- (DE3) *E. coli* strain was transformed with the expression plasmids, and protein expression was induced with 1 mM IPTG for 3 h at 37°C. GST and GST-fusion proteins were purified using glutathione sepharose beads (GE Healthcare, 17-0756-01) according to standard protocols. The purity of eluted and dialysed proteins was ascertained by SDS-PAGE, followed by staining with Coomassie Brilliant Blue R-250.

### Purification of FRAPPe and FRAPPe^Mut^

pRSET eCFP-CHAD-eYFP and pRSET eCFP-CHAD^Mut^-eYFP are transformed into separate BL21DE3 competent cells and incubated at 37℃ for 14 hours. A single colony of each from the agar plate was grown in LB medium with 100 μg/ml Ampicillin, and protein expression was induced with 1mM IPTG at 18℃ for 14 hours. Cells were pelleted by centrifugation at 6000 rcf at 4℃ for 20 min and then lysed by sonication (each cycle time- 2 min, output voltage 20, output control 30, duty cycle 30/40) till the lysate became less viscous in Tris-Cl pH 7.6 lysis buffer. The cell-free lysate was obtained by centrifugation at 10000 rcf at 4°C for 30 min. The supernatant was transferred to a fresh tube, and equilibrated Ni-NTA agarose beads were added.

This was incubated in a rotor at 4°C for 2 hours. The protein was then subjected to affinity chromatographic purification and eluted via gradient concentration of imidazole from 25mM to 200mM. Eluted protein fractions were concentrated and run through a Size Exclusion Chromatography column (Superdex200). The eluted protein of the desired size was concentrated, and the protein was quantified by Bradford assay. Single-use aliquots were flash-frozen in liquid nitrogen and immediately stored at −80°C.

Tris-Cl Lysis Buffer pH=7.6: 20mM Tris, 100mM NaCl, 10% glycerol Size exclusion and storage buffer pH=7.6: 20m Tris, 10% glycerol

### Isothermal Calorimetry (ITC)

The binding of FRAPPe to polyphosphate was assessed using PolyP 100 as the ligand. FRAPPe and PolyP 100 were prepared at final concentrations of 130 µM and 1.3 mM, respectively. A 100 mM stock solution of PolyP was prepared in deionised water and subsequently diluted into the dialysis buffer used for FRAPPe. ITC measurements were performed at 25 °C. FRAPPe was loaded into the sample cell, and PolyP 100 was placed in the injection syringe. The titration consisted of one initial 0.49 µL injection followed by fifteen 2.49 µL injections of PolyP 100 into the cell. After completing the FRAPPe titration, a reference experiment was performed by injecting PolyP 100 into the dialysis buffer under identical conditions. Heat signals from the buffer control were subtracted from the protein titration to obtain the final binding isotherm.

### Fluorescence Spectra measurements

With the help of the Tecan Infinite 200 Pro microplate reader, we measured the fluorescence intensity spectra of FRAPPe. To characterise FRET in the protein, we excited the protein at 415 nm wavelength and measured the fluorescence emission intensities across a spectrum ranging from 450 to 650 nm. The wavelength interval chosen for the readings was 1 nm. The protein concentrations used ranged from 200-4000 nM. The polyP∼50 concentrations used ranged from 0-25000 ug/ml. The protein and polyP∼50 dilutions are made with the protein storage buffer.

### Calculation of FRET ratio from fluorescence spectra

To calculate the FRET ratio, we took an average of FRET emission readings at 515-525 nm wavelength (520 nm maxima, F_a_) and divided it by the average of donor emission readings from 480-490 nm wavelength (485 nm maxima, F_d_). FRET ratio= F_a_/F_d,_ where F_a_ is the average of acceptor emission readings at 515-525 nm, and F_d_ is the average of donor emission readings at 480-490 nm.

### Negatively charged molecule solutions

We used negatively charged molecules, in addition to polyP∼50, to assess their effect on the sensor’s FRET activity. We used DNA, RNA, Heparin, monophosphate (Pi), dNTP, and pyrophosphate (PPi). The required dilutions for each molecule were made with Tris buffer (20mM Tris+10% glycerol), where effective concentrations in the assay were at 30 *μ*g/ml for DNA, 15 *μ*g/ml for RNA, 2.5 *μ*g/ml for Heparin, 1mM for Pi, 24*μ*M for dNTP, and 2.18*μ*M for PPi. PolyP∼50 concentrations ranged from 31250 *μ*g/ml to 0 *μ*g/ml with a total of 16 data points in the presence of all these ions to assess the difference in binding capacity of the sensor to polyP in the presence of the chosen negative polymers.

### FRET measurements

After setting up the reaction, we incubated the plate in the dark at room temperature for 10 minutes before measuring. The FRET measurements were taken using a BMG Labtech Polarstar Omega microplate reader with the excitation filter at 450 nm and dual emission filters at 485 nm and 520 nm, respectively. The fluorescence gain was set at 1000 for both emission filters. The reader’s temperature was set to 25 ℃ during the readings.

### FRET ratio and change calculation

The FRET emission measured at 520 nm is divided by the donor eCFP emission at 480 nm (=F_a_/F_d_). For the calculation of the FRET ratio change, the FRET ratio is calculated for the corresponding concentrations of the polyP∼50(F_n_) as well as for the blank, which has no polyP∼50 (F_0_). The difference between F_n_ and F_0_ with respect to F_0_ gives the FRET ratio change for that specific polyP∼50 concentration(=ΔF/F_0_).

FRET ratio= F_a_/F_d_= Em_520_/ Em_485_

FRET ratio change= 1- (F_n_/F_0_)= ΔF/F_0_

### Synthesis of Bis-biotin-PolyP_8_

Bis-biotin-PolyP_8_ was made as reported earlier (ACS Chem. Biol. 2018, 13, 8, 1958–1963). Briefly, bis-propargylamidate-PolyP_8_ was dissolved in triethylammonium acetate buffer (100 mM, pH 7). Azide-PEG3-biotin was added to the mixture, and subsequently, CuSO_4_ • 5 H_2_O and sodium ascorbate were added. The reaction mixture was stirred under argon for 3 h. Chelex® 100 was added, and the mixture was gently stirred for 15 min before filtering through a syringe filter. The crude product was precipitated by dropwise addition to a cold NaClO4 acetone solution, centrifuged, and washed with acetone three times. The dried crude product was purified by anion exchange chromatography (High Trap QFF, increasing concentration of aq. NaClO_4_-solution (1 M) in H_2_O). Fractions eluted with 7-10% aq. NaClO_4_ was combined, lyophilized, and precipitated with cold NaClO_4_ acetone solution (0.5 M). The blue solid was dissolved in water, and Chelex® 100 was added to remove the copper counterions. The mixture was stirred for 45 minutes at room temperature (r.t.) before being filtered through a syringe filter and precipitated with a cold NaClO_4_ acetone solution (0.5 M). This procedure was repeated till Bis-biotin-PolyP_8_ (MW: 1796.93) was obtained as a white solid. Its purity was determined using nuclear magnetic resonance (NMR) or capillary electrophoresis-mass spectrometry (CE-MS) and stored as an amorphous powder at 4°C or resuspended in nuclease-free water to store at −80°C.

### Binding of Bis-biotin-PolyP_8_ to GST-CHAD, GST-CHAD^Mut^ in a 96-well plate assay

Flat-bottom 96-well microtiter black plates (Sigma-Aldrich M4936) were used for the binding assay. The wells were coated with 100 ng of either GST, GST-CHAD wild-type or GST-CHAD^Mut^ in 50 µL bicarbonate buffer pH 9.0 (15 mM sodium carbonate, and 35 mM sodium bicarbonate) and incubated at 4°C for 12-16 h in a moist chamber. The wells were subsequently washed with Tris-buffer saline (TBS; 50 mM Tris pH 7.4, 150 mM NaCl), blocked with 2% BSA in TBS-T (TBS with 0.1% Tween 20) for 2 h at room temperature (RT), followed by three washes with TBS-T. Serial dilutions of decreasing concentration of bis-biotin PolyP_8_ were added to coated wells and incubated for 1 h at RT. Wells were washed three times with TBS-T, and 100 µL of streptavidin-HRP (Invitrogen, S911), diluted 1:10,000 in TBS, was added to the wells and incubated for 1 h at RT. Following three washes in TBS-T, 100 µL of luminol reagent SuperSignal^TM^ ELISA Pico Chemiluminescent substrate (Cat. No. 37070), diluted 1:5 in TBS, was added to each well, and incubated for 10 min at RT. Luminescence values in relative light units (RLU) were recorded using a multimode plate reader (Perkin Elmer 2300 EnSpire), and the buffer blank was subtracted from all readings. For saturation binding analysis, the specific binding of CHAD to each concentration of Bis-biotin-PolyP_8_ was calculated as follows: Specific binding = Total binding (GST-CHAD) - non-specific binding (GST). A specific binding curve was plotted, and Kd values were calculated using the one-site specific binding equation in GraphPad Prism 5. The binding of GST-CHAD^Mut^ to polyP was tested using Bis-biotin-PolyP_8_ (5 µM) in the absence or presence of PolyP_20_ (10^6^ pmoles of Pi), using the method described above. Data were compiled from 3-4 independent experiments. With this assay, we obtained a dissociation constant (Kd) of ∼1.5 µM for the binding of Bis-biotin-PolyP_8_ to CHAD **(Supplementary Fig. 2F)**. This is lower than the binding affinity reported for the interaction of *Ss*CHAD with biotinylated polyP_100_ (∼45 nM) using quantitative grating-coupled interferometry (29).

### Polyphosphate extraction

*Saccharomyces cerevisiae BY4741* (wild-type), and *vtc4Δ* (20 ml, OD=1.5) were lysed in LETS buffer (10 mM Tris-HCl, pH 8.0, 100 mM lithium chloride, 10 mM EDTA, 0.2% SDS); 200μl LETS buffer was added to the yeast pellet and 200μl glass beads. The mixture was centrifuged for 45 minutes at 4℃ for cell rupture and lysis. To precipitate the genomic DNA, acid phenol (pH 4.5), equivalent to the volume of the lysis buffer, was added, and the samples were centrifuged at 18000g for 10 minutes at room temperature. The aqueous phase, which is devoid of genomic DNA, was transferred to a fresh vial and mixed with a double volume of chloroform. The mixture was vortexed for three minutes and centrifuged at room temperature at 18000g for five minutes. The aqueous phase, devoid of proteins, was collected and treated with RNase A at 37℃ for four hours, followed by the addition of an equal volume of 1:1 phenol: chloroform mixture. The solution was vortexed for three minutes and centrifuged at room temperature at 18000g for five minutes. The aqueous phase was collected, and chloroform was added to it in a volume double that of the samples. The mixture was vortexed for three minutes and centrifuged at room temperature at 18000g for five minutes. The aqueous phase was collected and mixed with absolute ethanol in the ratio 1:2.5 (vol/vol), followed by overnight incubation at −80℃. The next day, the polyP was precipitated by centrifugation at 18000g at 4℃ for 30 minutes. Ethanol was decanted, and the transparent polyP pellet was air-dried before dissolving in water.

### Polyphosphate quantification using malachite green assay

Malachite green (MG) detects monophosphates released from polyP pools due to the addition of purified yeast exopolyphosphatase (*Sc*PpX1). Samples with extracted polyP were divided into two parts: one treated with 5 µg/ml ScPpX1 for 18 hours at 37°C, and the other served as the untreated control. MG reagent was prepared by mixing MG (0.045% in water) and Ammonium molybdate (4.2% in 4N HCl) in a 3:1(vol/vol) ratio and filtered through a Whatman grade 1 paper. K_2_HPO_4_ (0.1 – 20 nmoles) was used as a phosphate standard. Samples or standards (50 μl) were loaded into 96-well plates, and 200 μl of MG reagent was added per sample, followed by incubation in the dark for 15 minutes at room temperature. Absorbance was measured at 650 nm using the Omega PolarStar Plate Reader. PolyP content of the samples (in Pi terms) was determined by interpolation from a linear regression analysis of the K_2_HPO_4_ standard.

### PolyP extraction from mouse tissues

3 months old C57BL/6 J mice were sacrificed, the liver was harvested, and washed thoroughly with 1X TBS. Fresh weight of all the tissues was taken, and the tissues were minced. Homogenization was further carried out at a concentration of 250 mg of tissue per ml of 1X LET buffer with 2mM EGTA. To 1 ml of tissue homogenate, 1 ml of acidic phenol was added and mixed by vortexing for 5 seconds. The mixture was then centrifuged for 10 min at 18,000 g. The aqueous phase was then transferred to a fresh tube, and an equal volume of chloroform was added, followed by vortexing and a second spin at 18,000 g for 10 min. To the supernatant, 2.5 times absolute ethanol was added and kept at −80°C overnight. A high-speed spin (21,000 g) for 30 minutes at 4 °C was then given to the samples, resulting in polyP pellet formation. The supernatant was discarded, the pellet was air-dried, and the pellet was resuspended in 110 µl of HPLC water. 100 µl of the suspension was used for the FRAPPé assay, while a 10 µl aliquot was further used for the DAPI assay.

### Polyphosphate quantification using DAPI assay

DAPI assay for the measurement of polyP from tissue was carried out following the protocol of Ladke et al. (36). The 10 µl aliquot of polyP pellet suspension was mixed with 10 µl of 10X recording buffer and 80 µl HPLC water. It was divided into two tubes with 50 µl of sample in each. The sample in one tube was treated with ScPPX (1 µg), and another tube was kept untreated. Both treated and untreated tubes were incubated overnight at 37 °C. For the assay, a standard curve was prepared using a 2X dilution of SHP (polyP20) in the recording buffer, ranging from 10,000 pmoles to 300 pmoles. In a 96-well fluorescence microplate, 100 µl of standard and 25 µl of each treated and untreated sample were added in duplicates. A recording buffer was added to the samples to make up the final volume to 100 µl. To each sample and standard, 100 µl of DAPI (60 µM in recording buffer) was added, and the plate was kept in the dark for 15 min. The fluorescence was then measured using excitation of 415 nm, and emission was collected at 550 nm using a Perkin Elmer EnSpire multimode plate reader. The difference between the interpolated values for the treated and untreated samples was calculated to yield the polyP concentration. Recording buffer: 20mM HEPES pH-7.4, 150mM KCl.

### Graph and statistical analysis

All the graphs and statistical analysis were done using GraphPad Prism software. The statistical tests used are mentioned in the respective figure legends.

### Materials and reagents List

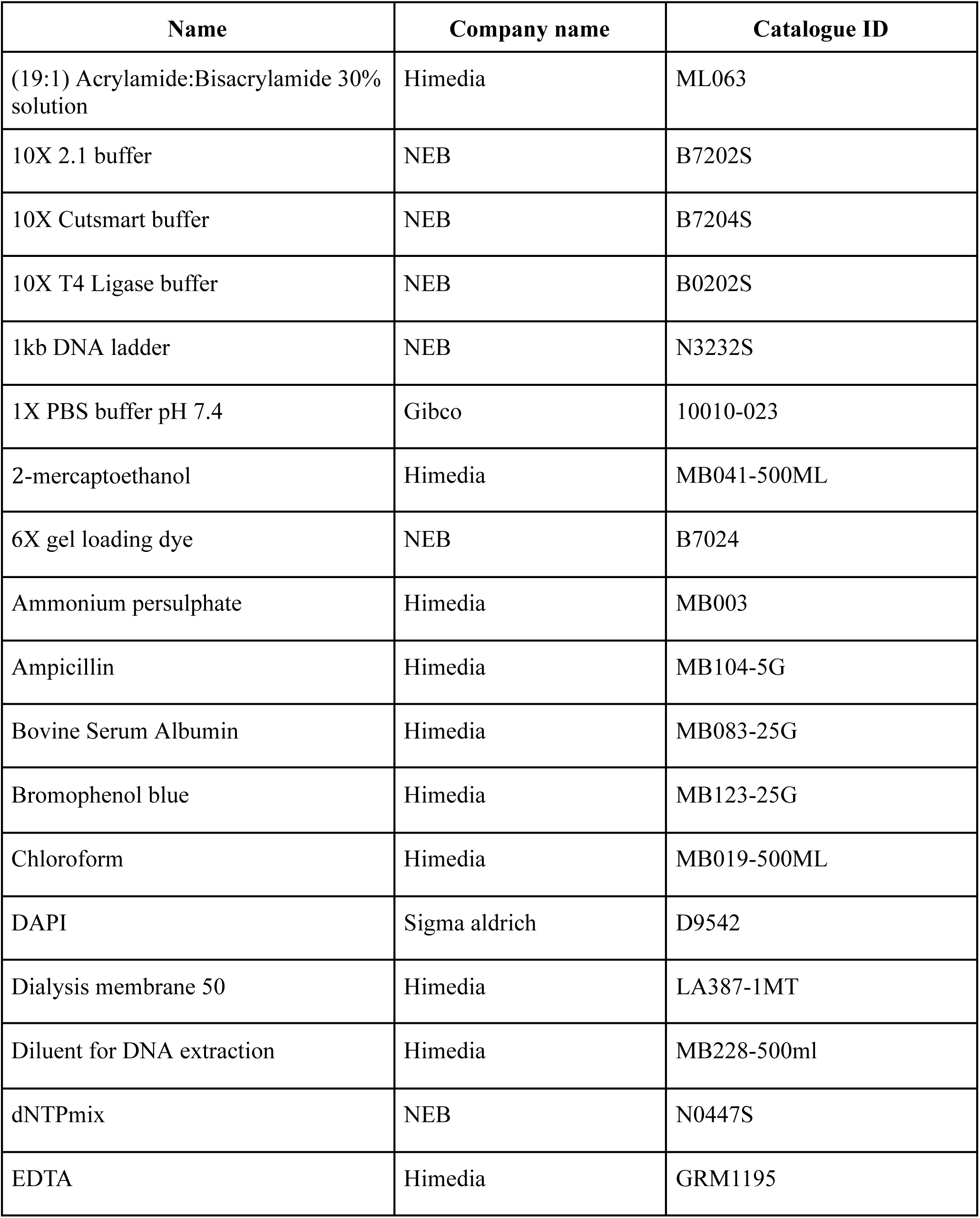
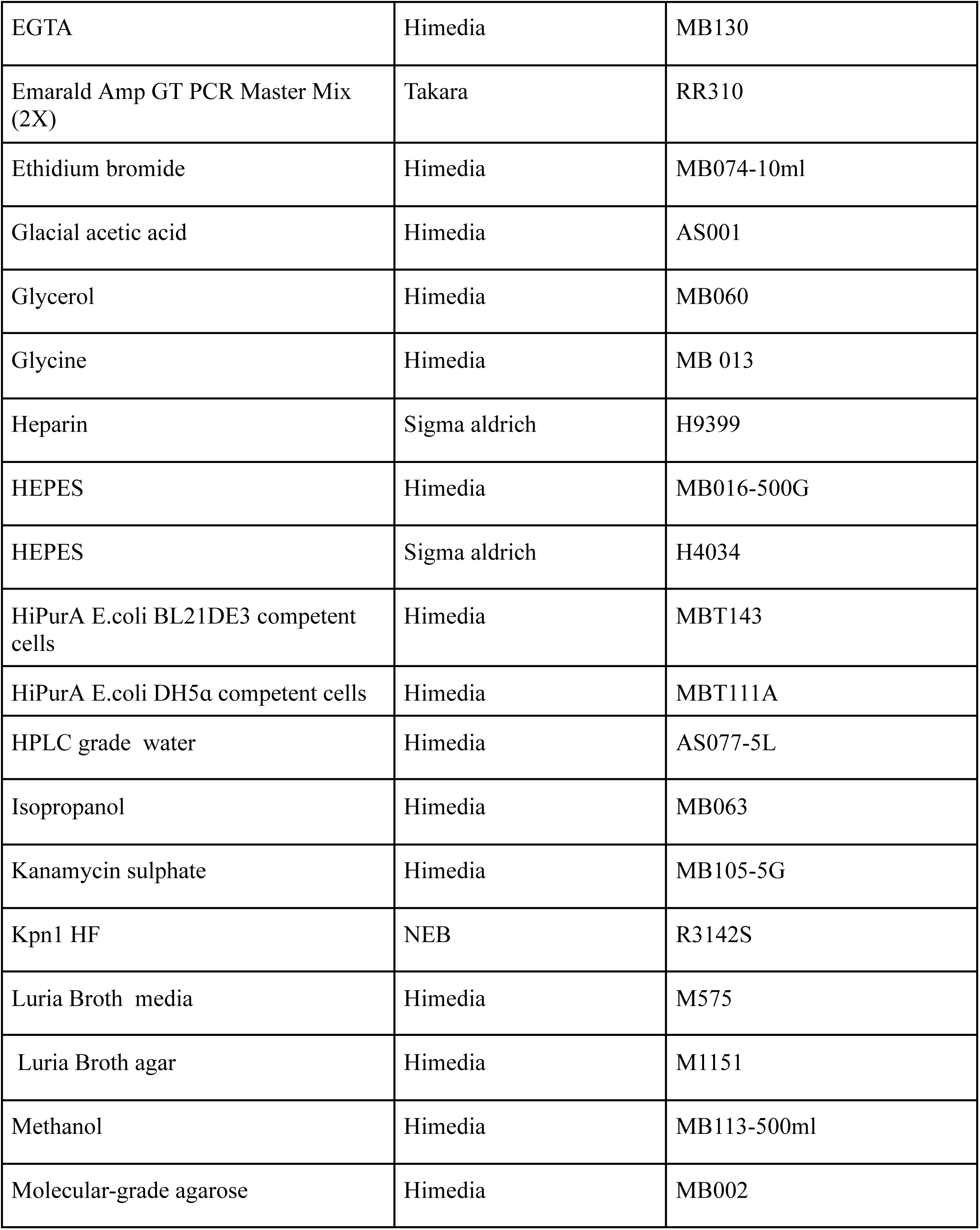
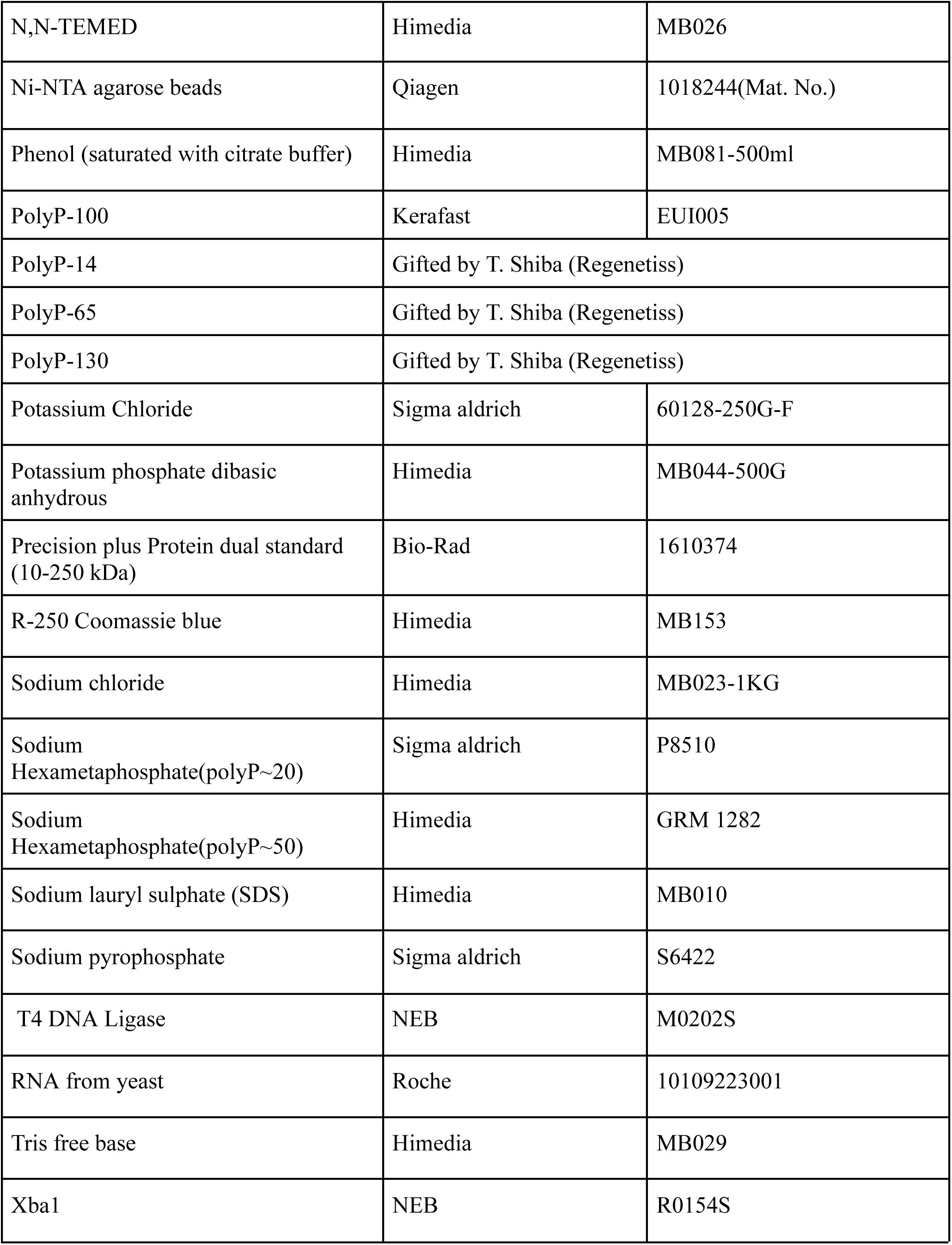
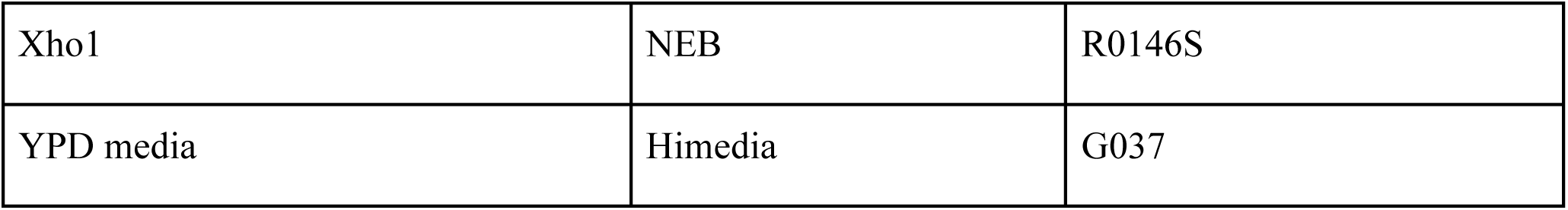

### Plasmid List

| Name | Company name | Catalogue ID |
| --- | --- | --- |
| pMH-HT_SsCHAD | addgene | 127783 |
| pRSET FLIPPi 260n | addgene | 13552 |
| pGW CHAD | Genewiz | Synthesised |
| pGW CHAD<sup>Mut</sup> | Genewiz | Synthesised |

### Primer List

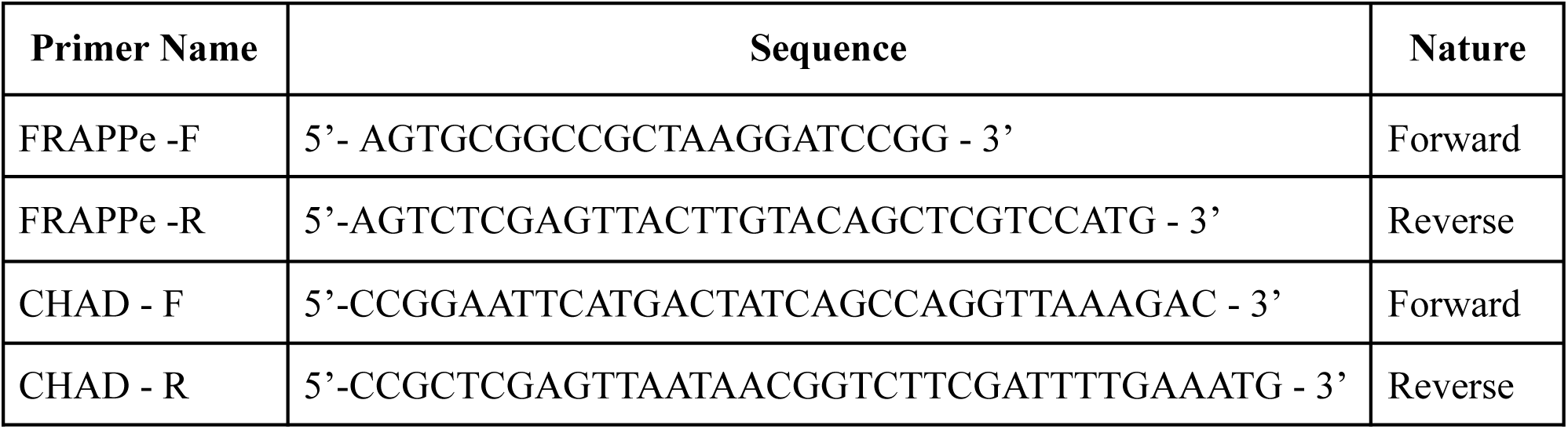

### Kit List

| Name | Company name | Catalogue ID |
| --- | --- | --- |
| Gel extraction and PCR cleanup kit | Machary Nagel | 740609.25 |
| HiPurA E.coli plasmid miniprep kit | Himedia | MB508 |
| Pierce detergent-compatible Bradford assay kit | Thermofisher | 23246 |

## Acknowledgements

We thank T. Shiba for generously sharing polyP of different chain lengths -polyP (PolyP_14_, PolyP_65_, PolyP_130_). We thank Anand T Vaidya and Vipin Agrawal for their advice and support for protein purification. We thank Debaditya De, Vineesha Oddi, Akruti Shah from Centre for DNA Fingerprinting and Diagnostics (CDFD), India, and Jyoti Singh from University of Freiburg, Germany, for their contributions and assistance in the Bis-Biotin PolyP_8_ synthesis and the 96-well plate Bis-Biotin PolyP_8_ binding assay. We thank Kalyaneswar Mandal and members of our lab for discussions and valuable feedback. We thank Aneesh T. Veetil for the critical reading, comments, and suggestions on the manuscript.

## Funding

This work is funded by an Indo-German joint research grant by the Department of Biotechnology, New Delhi, India (IC12025(11)/2/2020-ICD-DBT) and Deutsche Forschungsgemeinschaft (DFG, German Research Foundation, project number 445698446) to MJ, RB, and HJJ. MJ acknowledges support from the Department of Atomic Energy (Project Identification No. RTI4007), Department of Science and Technology, SERB (CRG/2020/003275), Department of Biotechnology (BT/PR32873/BRB/10/1850/2020), Government of India. RB acknowledges support from the Department of Biotechnology, Ministry of Science and Technology, Govt. of India (BT/PR29960/BRB/10/1762/2019); Science and Engineering Research Board, Department of Science and Technology, Govt. of India (CRG/2019/002597); and CDFD core funds. AK is supported by the M. K. Bhan Young Researcher Fellowship, Department of Biotechnology, Government of India (HRD-16016/1/2025-HRD-DBT E-20859). Moreover, this work was supported by the Deutsche Forschungsgemeinschaft (DFG) under Germany’s Excellence Strategy (CIBSS-EXC-2189-Project ID 390939984, to HJJ). SS received travel support from TIFR-Infosys Leading Edge Travel Grant (Leading Edge TG/(R-11)/09/), and Anusandhan National Research Foundation- International Travel Support (ANRF-ITS) (ITS/2025/000980).

## Author contributions

SS, MJ, and VB have conceptualised the study and designed the FREPPe. SS: Designed the methods, did cloning and protein purification, performed experiments, validated, and analysed the data. PA and HS have cloned various FRAPPe constructs. PA, DRP, and SS: Purified FRAPPe and performed all FRET experiments. AK and RB: Mouse polyP extraction and quantification. SRK: Performed the ITC experiments. SM and HJ: Synthesized the Bis-biotin PolyP_8_. MM and RB: Purified GST, GST-CHAD, and GST-CHAD^Mut^ proteins and performed Bis-biotin PolyP_8_ binding assay. VB and KRM contributed to FRET data analysis and guided experimental designs. MJ supervised the study and was responsible for the project administration and funding acquisition. SS and MJ wrote the original draft. Everyone has reviewed and edited the manuscript.

---

**Supplementary Fig. 1:**
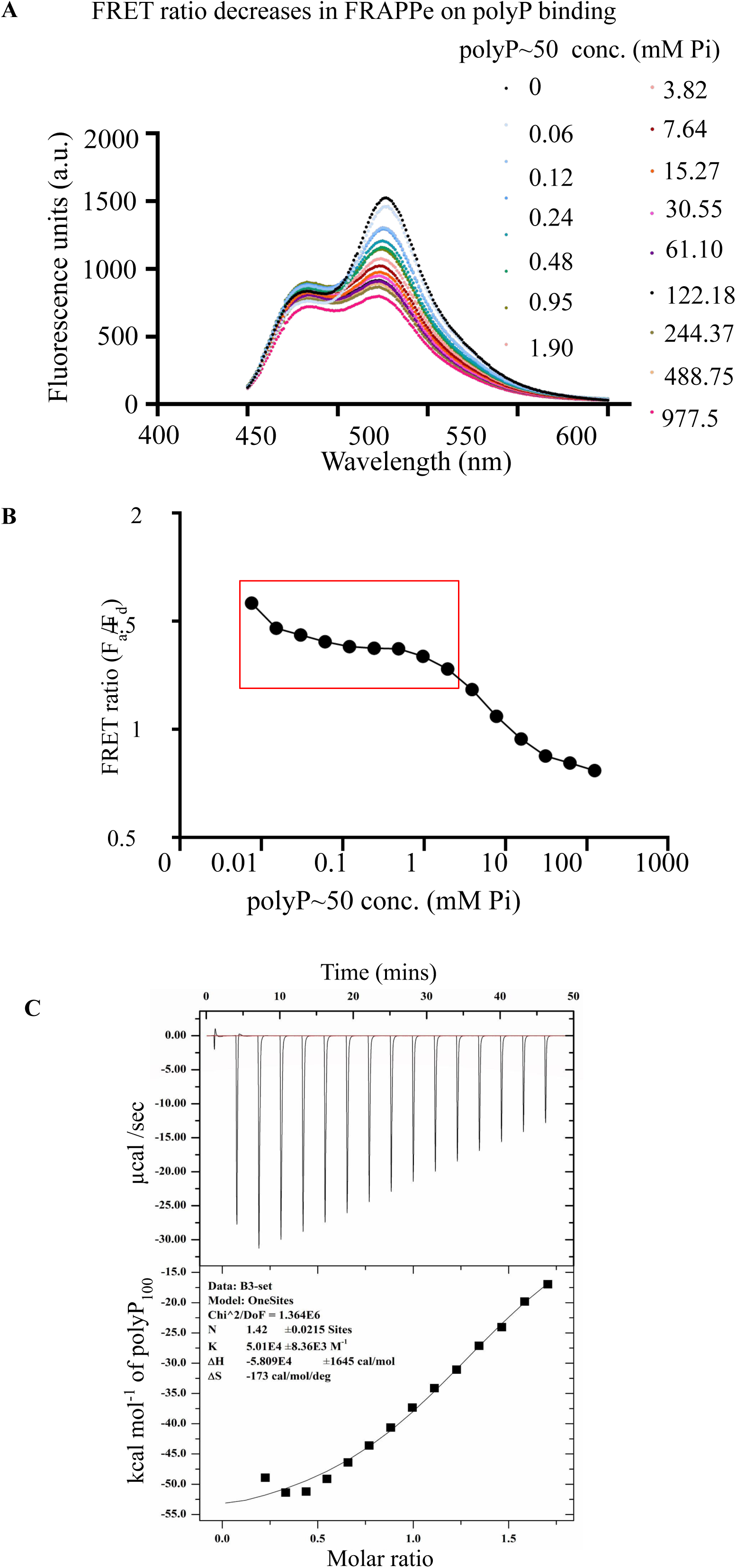
Characterisation of FRAPPe. **A)** Fluorescence spectral scan (450nm-650nm) of 0.75 µM FRAPPe on excitation at 415 nm, with varying concentrations of commercially available polyP∼50. **B)** The FRET ratio calculated from *1A* shows a two-step binding characteristic curve. The red box highlights the range used for subsequent experiments, as it corresponds to physiological polyP concentrations reported in mammals. **C)** Representative ITC binding isotherm for the interaction between PolyP-100 and FRAPPe in 20 mM Tris, pH 7.4. Fitting of the isotherm yields an apparent binding stoichiometry (N) of 1.42.

**Supplementary Fig. 2:**
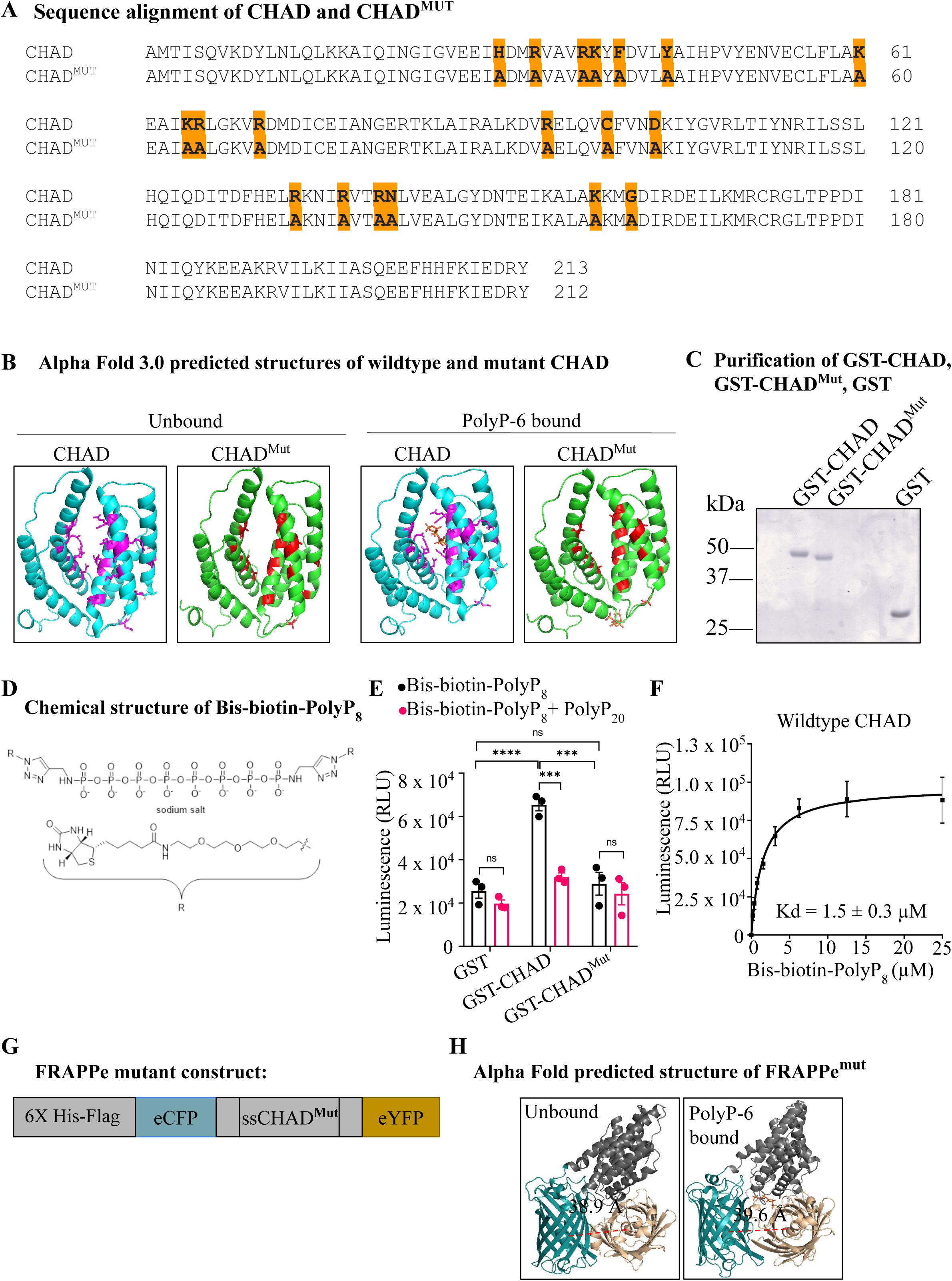
Characterisation of CHAD and CHAD^Mut^. **A)** Sequence alignment of CHAD and CHAD^Mut^, highlighting the amino acids that have been mutated to Alanine to destroy polyP binding. **B)** Alpha Fold 3.0 predicted structures of CHAD and CHAD^Mut^, unbound and bound to polyP-6 (docked), showing that CHAD binds to polyP but CHAD^Mut^ loses the polyP binding at the binding pocket. **C)** Chemical structure of Bis-biotin-PolyP_8_, where R indicates PEG3-biotin. **D)** Bacterially expressed and purified GST, GST-CHAD, and GST-CHAD^Mut^ resolved by SDS-PAGE, and visualised by staining with Coomassie Blue R 250. **E)** Saturation binding curve for specific binding of Bis-biotin-PolyP_8_ to immobilised GST-CHAD (mean ± SEM, N=4). Data was analysed in GraphPad Prism 5 using the one-site specific binding equation after subtracting non-specific binding to GST. **F)** Binding of GST, GST-CHAD wild-type, and GST-CHAD^Mut^ to Bis-biotin-PolyP_8_ in the absence and presence of competing PolyP_20_ (mean ± SEM, N=3). The data were analysed using two-way ANOVA with Tukey’s multiple comparisons test (****P<0.0001; ***P≤0.001; ns, not significant, P>0.05). **G)** Schematic of the FRAPPe^Mut^ construct, and **H)** Alpha Fold 3.0 predicted model of FRAPPe^Mut^ (unbound) and polyP-6 docked (bound).

**Supplementary Fig. 3:**
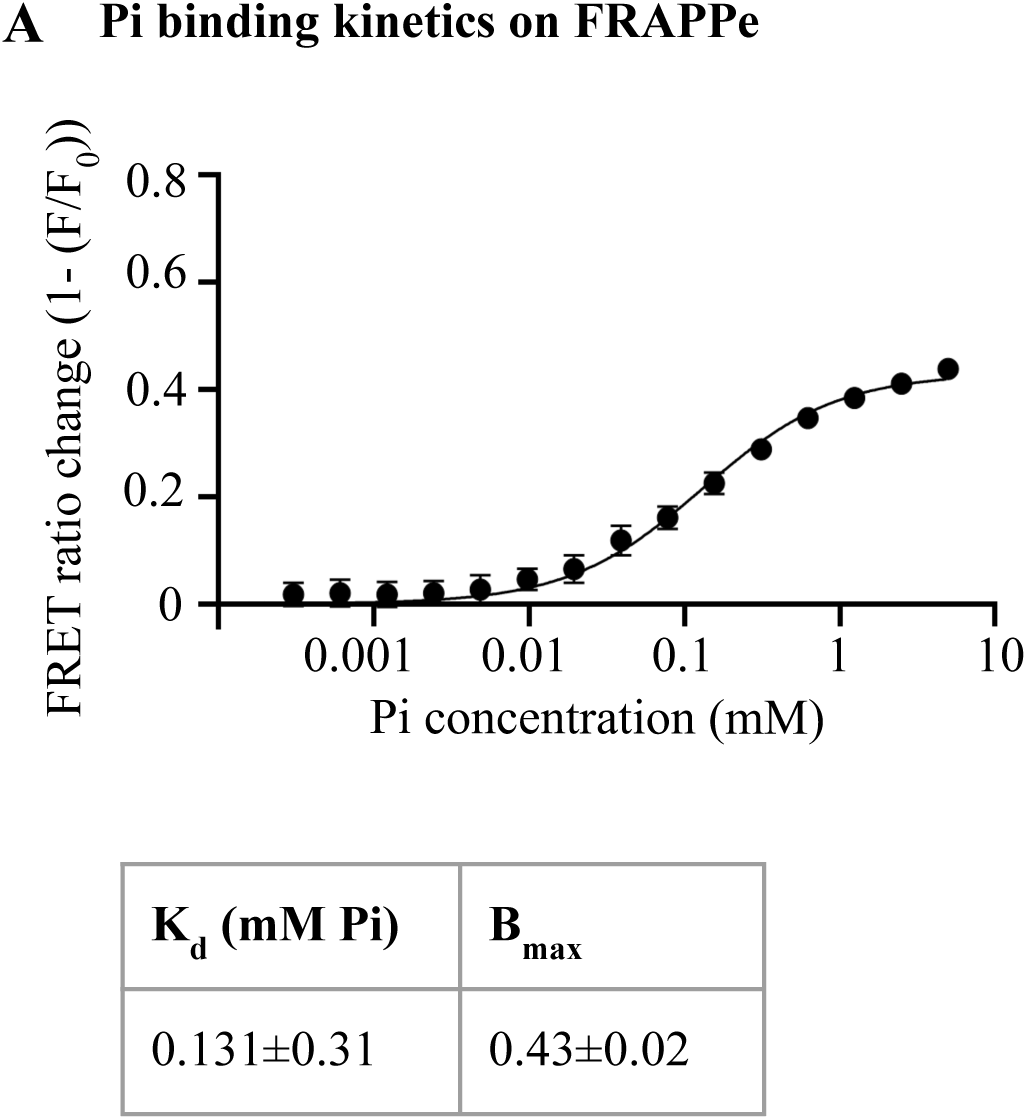
**A)** FRET ratio change of FRAPPe across KH_2_PO_4_ (Pi) concentration range 0-5 mM in 20 mM Tris buffer. The chart represents the binding affinity (K_d_) and the binding capacity (B_max_) of FRAPPe towards monophosphates. Data plotted with s.e.m. as an error bar.

## Notes

### Competing Interest Statement

The authors have declared no competing interest.

